# Genome-wide reconstruction of complex structural variants using read clouds

**DOI:** 10.1101/074518

**Authors:** Noah Spies, Ziming Weng, Alex Bishara, Jennifer McDaniel, David Catoe, Justin M. Zook, Marc Salit, Robert B. West, Serafim Batzoglou, Arend Sidow

## Abstract

Recently developed methods that utilize partitioning of long genomic DNA fragments, and barcoding of shorter fragments derived from them, have succeeded in retaining long-range information in short sequencing reads. These so-called read cloud approaches represent a powerful, accurate, and cost-effective alternative to single-molecule long-read sequencing. We developed software, GROC-SVs, that takes advantage of read clouds for structural variant detection and assembly. We apply the method to two 10x Genomics data sets, one chromothriptic sarcoma with several spatially separated samples, and one breast cancer cell line, all Illumina-sequenced to high coverage. Comparison to short-fragment data from the same samples, and validation by mate-pair data from a subset of the sarcoma samples, demonstrate substantial improvement in specificity of breakpoint detection compared to short-fragment sequencing, at comparable sensitivity, and vice versa. The embedded longrange information also facilitates sequence assembly of a large fraction of the breakpoints; importantly, consecutive breakpoints that are closer than the average length of the input DNA molecules can be assembled together and their order and arrangement reconstructed, with some events exhibiting remarkable complexity. These features facilitated an analysis of the structural evolution of the sarcoma. In the chromothripsis, rearrangements occurred before copy number amplifications, and using the phylogenetic tree built from point mutation data we show that single nucleotide variants and structural variants are not correlated. We predict significant future advances in structural variant science using 10x data analyzed with GROC-SVs and other read cloud-specific methods.

## Introduction

Structural variants (SVs) represent the highly heterogeneous class of large-scale changes in the genome, encompassing DNA edits that include but are not limited to deletions, tandem duplications, inversions, translocations, and combinations that are generally referred to as 'complex events'. Because each event affects a large genomic region, SVs are responsible for the majority of nucleotides varying between individuals^1^ and in many cancer genomes^2,3^.

Despite its importance in evolution and disease, structural variation remains difficult to comprehensively characterize. DNA breakage and subsequent fusions can connect any genomic locus to any other, and therefore the potential search space for variant detection is proportional to the square of the genome size. (The search space for small variants is proportional only to the genome size.) Repetitive loci, uneven or biased sequencing coverage, and the typically short length of sequenced fragments complicate accurate detection. Thus, for example, while the presence of a large duplication may be easy to identify from an increase in sequencing coverage, the exact breakpoints and location of the duplicate copy may be undetectable in data from current sequencing methods. Copy number-invariant types of structural variation, such as inversions and translocations, or smaller copy-number altering variants, are also difficult to detect and characterize.

Previous work has illuminated the potential complexity of SVs^4–7^. One example of large-scale complexity is chromothripsis^3^, in which a chromosome shatters into many pieces that are then apparently randomly reassembled, leading to massive rearrangements and loss of heterozygosity in the intervening sequences. SVs may also exhibit local complexity arising from error-prone repair mechanisms that can, for example, result in insertion of short sequences at the sites of larger deletions (reviewed in ref. 8).These complex events can be difficult to interpret using existing sequencing technologies. For example, analyses of short-fragment sequence data can only confidently relate breakpoints that are within the fragment size distribution, typically <500 bp. Longer-distance reconstruction (e.g. ref 2) requires the assumptions that downstream events occur in the same haplotype and that all breakpoints have been accurately identified. Single-molecule long-read approaches are better suited for detection of SVs, but throughput and cost are typically limiting, and the high per-base error rate is a drawback.

An alternative to long reads is short-read sequencing of bits of originally long fragments, as in mate-pair libraries, where both ends of a long fragment are brought together into a short fragment that can be Illumina-sequenced^9^. The resulting data exhibit very high physical coverage relative to sequence coverage such that a single-copy SV breakpoint is typically covered by hundreds of mate-pairs^10^. However, high-quality mate-pair libraries are difficult to generate and are practically limited in their fragment lengths.

Read clouds represent the next generation of the long-fragment / short sequence approach, marrying the advantages of standard Illumina sequencing (high throughput and accuracy) with long-fragment information added through a barcode tag incorporated during a molecular partitioning step^11–13^. We have previously shown that such an approach can improve the mapping of short reads to repetitive genomic regions^14^. The recently released 10x Genomics platform produces read cloud libraries with dramatically higher numbers of partitions compared to previous methods, enabling new applications^15^.

To prepare 10x Genomics libraries, long DNA fragments are diluted into ~10^5^ (previous generation, *GemCode*) to 10^6^ (current generation, *Chromium*) microfluidic droplets, each of which contains a unique barcode. Within each droplet, randomly primed amplification produces many short fragments templated off the long fragments, all of which share that same barcode. When these barcoded short fragments are Illumina-sequenced, their alignments to the reference genome form clusters. We refer to the clusters of identically barcoded, linked reads, as clouds. Each individual cloud of linked reads is a sample of sequence information from the originating long fragments.

The long-range information in read clouds can in principle be leveraged to identify, sequence-assemble, and reconstruct complex SVs. Using a novel method that we developed for this purpose, **G**enome-wide **R**econstruction **o**f **C**omplex **S**tructural **V**ariants (GROC-SVs), we show that 10x data substantially improves detection of SVs compared to standard short-fragment sequencing and that it enables the reconstruction of much more distant events compared to mate-pair sequencing. In addition, we use the read cloud information to produce high-quality assemblies of the sequences spanning the breakpoints, enabling us to better interpret local complexity. We applied GROC-SVs to characterize chromothripsis and subsequent evolution of structural variation in a liposarcoma and to analyze SVs in a breast cancer cell line.

## Results

### Sequence Data Generation and Characteristics

We Illumina-sequenced 10x GemCode libraries from each of 7 spatially distinct sites within a well-differentiated liposarcoma, as well as a matched control sample from the kidney of the same patient. For purposes of comparison and validation, we also sequenced PCR-free Illumina libraries from all 8 samples to ~35x sequence coverage, and, from 3 of the 7 sites plus the control, long-insert (~7kb) mate-pair libraries to ~20x sequence coverage.

We size-selected the sarcoma DNAs prior to 10x library preparation, resulting in a tight fragment size distribution (mean > 30 kb, depending on sample; 95^th^ percentile = ~80kb; Supplementary Fig. 1), with half of the bases in fragments longer than 53 kb (N50). After filtering, the libraries had ~170,000 barcodes per sample. We estimated coverage of the genome by long fragments, C_F_ (physical coverage, see ref. 14), to be ~250x. Coverage of each long fragment by short reads, C_R_, was ~0.10x, meaning that an average of 10% of positions in each long fragment were covered by reads. Thus, the overall sequence coverage per sample was C = C_R_ x C_F_ = 25x.

In addition to the liposarcoma case, we also analyzed Chromium data from the HCC1143 breast cancer and matched-normal cell lines. Prepared without size-selection, HCC1143 fragment sizes covered a wide distribution (mean=41 kb; 95^th^ percentile = 148kb; N50 > 80 kb). After filtering, there were ~700,000 barcodes per cell line. Physical coverage was C_F_=145x, and sequence coverage of each fragment was C_R_=0.34x, resulting in an approximate overall sequence coverage of 49x.

### Overview of GROC-SVs

We developed new methods to leverage the long-fragment information inherent in 10x data for the purpose of SV identification and characterization. We begin by looking for statistical evidence of inferred long fragments that span breakpoints. This is accomplished by quantification of barcode similarity between all pairs of genomic locations (Figure 1a). Levels of barcode similarity are highest between any two nearby loci since input long fragments tend to overlap both loci (Figure 1a, *diagonal*). Loci separated by distances larger than the input fragment size share zero or only a small number of barcodes. (This is because each barcoded partition contains only a small number of fragments randomly drawn from the genome, and thus the chance that multiple partitions contain long fragments from the same two distant loci is small; Figure 1a, *background*). Thus, the presence of multiple barcodes that are shared between two distant locations at a level higher than that background is indicative of a breakpoint where the two locations are joined (Figure 1a, *translocation*). Subsequent to breakpoint identification and refinement we perform sequence assembly of the linked reads from the relevant breakpoints. This includes the reconstruction of complex events on the basis of breakpoints that are connected by long clouds (Figure 1b).

**Figure 1.**
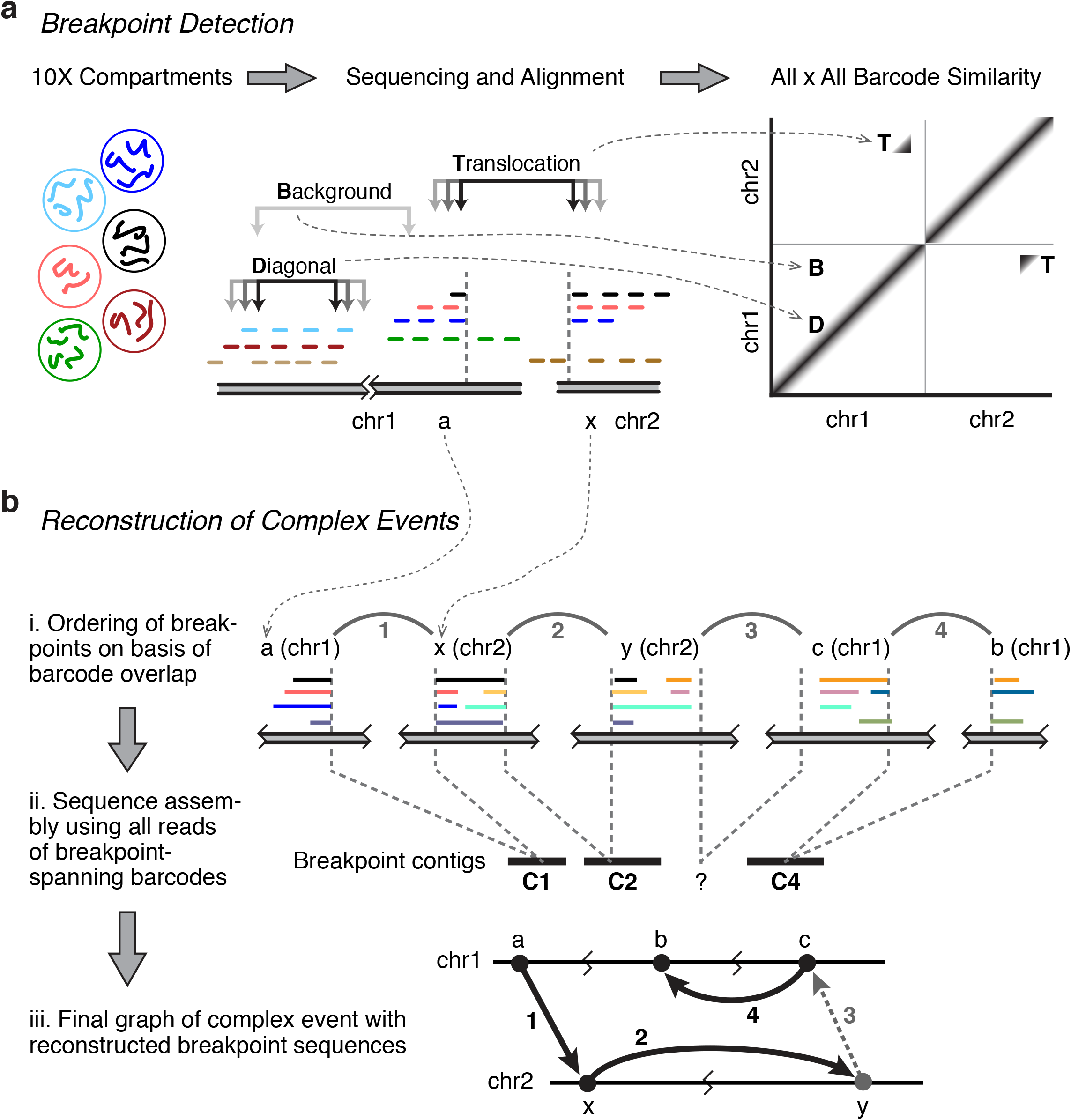
Overview of GROC-SVs. (a), explanation of the origin of the barcode similarity signal. 10x compartments are barcoded, signified by color; reads (short colored lines) linked by same barcodes form clouds upon alignment to the reference genome. Black, orange, and blue clouds span a translocation breakpoint that produces a characteristic signal off the diagonal; green and light brown originate from the other allele. Cyan, dark brown, and beige are not involved and are shown to illustrate the signal emanating from the pairwise comparison of nearby coordinates. (b), breakpoint graph construction and sequence assembly of complex events. Letters (a, b, c, for chromosome 1; x, y for chromosome 2) indicate genomic segments, numbers are the breakpoint connections, in order. Breakpoint 3 illustrates that not all high-confidence breakpoints yield an interpretable sequence assembly, but that they are still part of the reconstruction.

### Structural variant discovery with GROC-SVs: breakpoint detection

Leveraging the long-fragment information embedded in 10x data to identify breakpoints begins by quantifying barcode similarity between all pairs of genomic regions. Barcode similarity is highest near the breakpoint, and drops off from either breakpoint at distances proportional to the fragment size distribution (Figure 2a; see Supplementary Fig 2 for a more detailed explanation). Some independent fragments with the same barcode can cause a low level of background similarity, typically <1 (Chromium) or 0–5 (GemCode) barcodes at any given pair of positions. Corresponding barcode similarity in the matched normal sample is within the expected range for background (Figure 2b).

**Figure 2.**
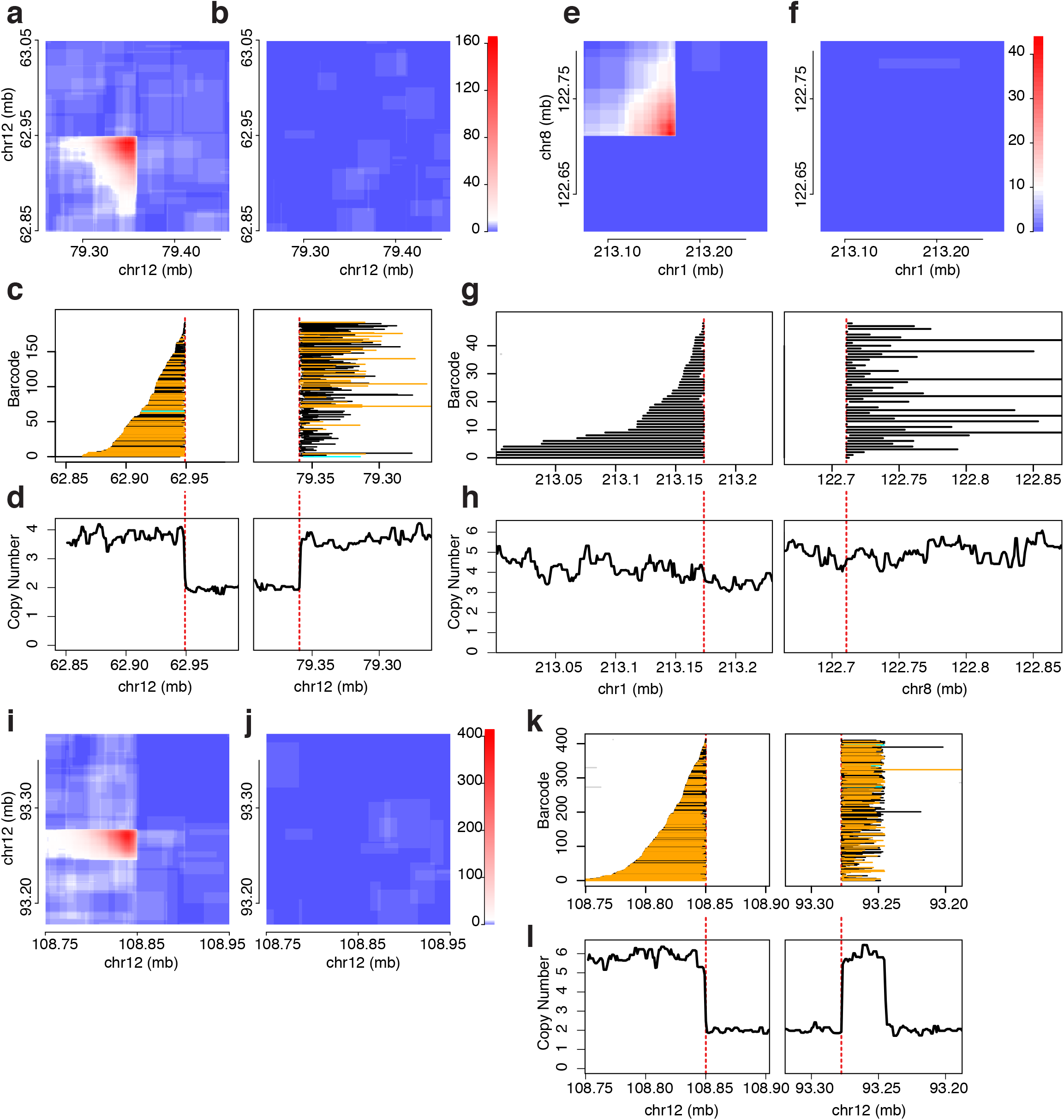
Examples of breakpoint signals in 10x data. (a – d), a simple breakpoint in sarcoma sample 0, Gemcode data. (e – h), a simple breakpoint in breast tumor cell line HCC1143, Chromium data. (i – l), two breakpoints in close proximity in sarcoma sample 0. (a, e, i), barcode similarity histograms in tumor. For each pair of genomic locations, the number of shared barcodes is color-coded according to the scale on the right, with the greatest signal forming a corner shape whose point is at the breakpoint coordinates. (b, f, j), same locations in the control samples. (c, g, k), inferred extent of breakpoint-supporting read clouds (corresponding to input fragments). Each row is one cloud, colored according to its assignment to a haplotype: supporting haplotype, orange; unassigned, black; non-supporting cloud in the same barcode as a supporting cloud, grey. Ordering is the same in left and right panels, revealing the difference between size-selected input DNA fragments from the sarcoma (c) and the broader distribution of unselected fragments from the cell line (g). (d, h, l), copy number profiles based on the short fragment data in the sarcoma (d, l), where a doubling of copy number is associated with the SV, or on the 10x data of the cell line (h), where no copy number change is evident. Decreasing coordinates indicate depiction of minus strand.

All supporting read clouds end near the putative breakpoint location (Figure 2c), a signal that is used during breakpoint refinement. In size-selected samples (as in the sarcoma) the clouds, ordered by their position relative to the first side of the breakpoint, tile across the breakpoint such that those starting furthest from the breakpoint tend to extend the least into the second region, while those starting closest to the first side extend the furthest into the second. Short-fragment sequencing coverage profiles support changes in copy number at many structural variant breakpoints (Figure 2d).

Barcode similarity is lower for a typical translocation from the HCC1143 cell line, presumably due to the lower physical coverage or differences in copy-number of the event (Figure 2e–h). However, because the Chromium data has more partitions, there are fewer independent fragments per barcode and the background is substantially lower. Thus, we observe essentially no background in irrelevant regions in the tumor data (Figure 2e) or anywhere in the corresponding regions of the matched normal cell line data (Figure 2f).

A third example, also from the sarcoma, illustrates the nature of the barcode similarity when two breakpoints are in close proximity. It shows the expected sudden dropoff in signal at the 108.85 mb breakpoint, but along the Y axis the signal ends abruptly not only at 93.27 Mb but also, in the other direction, at 93.25 Mb (Figure 2i; the control only exhibits background, Figure 2j). When tiling the read clouds, it becomes apparent that there are two breakpoints present at the 93 mb locus (Figure 2k), with copy number profiles exhibiting consistent levels that change abruptly at the breakpoint locations (Figure 2l). A substantial number of fragments appear to span from the first to the second breakpoint, suggesting that it is possible to use the 10x long-fragment information to directly link breakpoints that are in proximity to one another (see below).

In addition to providing high physical coverage of structural variant breakpoints, the long-fragment information in the 10x data allows for phasing of small variants with respect to the germline haplotypes^15^. Read clouds overlapping a heterozygous short variant can be assigned to one of the haplotypes. The low sequence coverage C_R_ of each fragment means that some read clouds, especially shorter ones, will not cover a short variant informative for haplotype assignment. However, the high physical coverage C_F_ results in a high total number of phased fragments for most genomic regions.

Because the structural variant breakpoints are distant from one another in the genome, the haplotypes are called independently for each side of the breakpoint, and so the standard phasing process does not uncover the phase arrangement for the tumor genome. However, nearly all informative fragments near each breakpoint support a single haplotype indicating that each side of the breakpoint only contributes a single haplotype to the event (Figure 2c,k). Thus we can use the predominant haplotype on either side of a breakpoint to locally phase the genomic regions that participate in the SV.

### Structural variant discovery with GROC-SVs: Sequence assembly of breakpoints and reconstruction of complex events

To better characterize breakpoints, GROC-SVs attempts to perform sequence assembly. We use the barcode information to identify relevant short reads that are fed into the assembler.

First, we identify barcodes that are shared among multiple breakpoints, suggesting some long fragments spanned across them; breakpoints that do not share barcodes are retained as singletons. For each such event or collection of events, we identify barcodes supporting each breakpoint and gather all reads marked by those barcodes (Figure 1), including those that were unmappable or had low mapping quality in the initial genome-wide mapping.

We then perform sequence assembly on these reads. As each barcode marks multiple fragments, many of the reads do not derive from a breakpoint-supporting genomic region. However, because fragments are randomly assigned to a barcode, these non-supporting fragments should be distributed randomly throughout the genome. Thus, combined sequence coverage is highest near the breakpoints, which should be covered by every barcode, and low elsewhere. Therefore, most assembled contigs actually derive from the SV haplotype. The assembled contigs are then aligned against the reference genome and those that support an SV are used to identify the exact breakpoint location.

The sarcoma had 12 events with 4 or more breakpoints, and 60 events with 2 or 3 breakpoints. As a fraction of all somatic breakpoints, 204/503 (41%) were assigned to complex events made up of at least 2 breakpoints. The ordering and assembly of 5 breakpoints comprising a sample complex event that spans 75 kb (Figure 3a–c and Supplementary Fig. 3) illustrates how the clouds tile and thereby connect neighboring breakpoints. Internal segments that could not be phased due to their short length are phased by virtue of being part of longer clouds that span breakpoints (Figure 3a). Copy number profiles are consistent with the reconstruction and reveal a subsequent 4-fold fold amplification of the variant (Figure 3b). Strikingly, the variant connects sequence from all over the long arm of chromosome 12 (Figure 3c).

**Figure 3.**
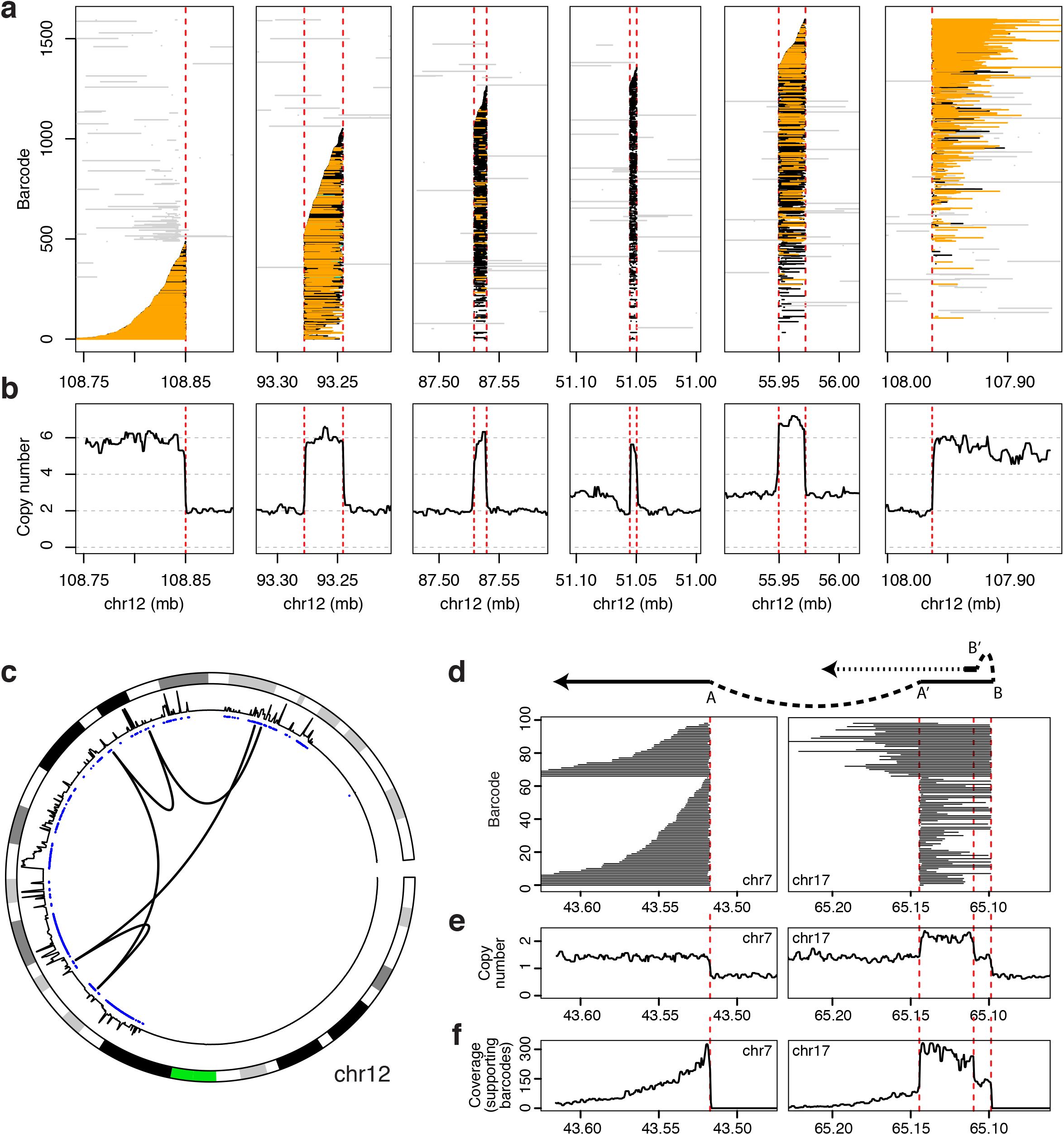
Reconstruction of complex events. Read clouds that support a complex event in the sarcoma. Clouds, colored as in Figure 2, tile across 5 consecutive breakpoints (a) with consistent copy number profiles (b). Circos plot with arcs depicting breakpoint connections illustrates that the event connects distant segments from the long arm of chromosome 12 (c). From outside to inside, chromosome ideogram (orange indicates the location of the centromere), then copy number profiles, then copy number aberration calls (blue for amplifications, red for deletions) are shown. A complex event in cell line HCC1143 (d) and its corresponding sequence read coverage (e, f).

In the breast cancer cell line, which did not undergo chromothripsis, we reconstructed 11 complex somatic events with a total of 24 breakpoints, including one event that illustrates both the potential complexity of structural variation and the power of read clouds to resolve it (Figure 3d–e). The sequence assembly, initiated by a breakpoint linking chromosomes 7 and 17, identifies a second breakpoint downstream on chromosome 17 involving an inversion, skipping approximately 10 kb of sequence and then resulting in a duplication of the downstream sequence (Figure 3d). Sequence coverage profiles show changes at the breakpoints (Figure 3e) that, upon analysis of only those reads that belong to the phased clouds, reveal the duplication. Without the long fragment information, it would have been impossible to show that the translocation and the inverted repeat were in the same tumor haplotype, nor could we have concluded that the inverted repeat continues beyond the translocation breakpoint.

### Genome-wide SV discovery, comparison and validation

The sarcoma genome harbored substantial structural variation, represented by a total of 503 called somatic breakpoints (Figure 4a). The vast majority fell within or at the edges of copy number amplifications, typically of relatively short genomic segments. The highest density of events occurred on the long arm of chromosome 12, involving 174 breakpoints (Figure 4a). Many events were subsequently amplified to high copy number, which exhibited high concordance between the two sides of the breakpoint (Figure 4b). These results provide further evidence for the recently described mechanism, also from a liposarcoma and also involving chromosome 12, by which chromothripsis is followed by breakage-fusion-bridge amplifications of neochromosomes, resulting in very high-copy number, rearranged genomic segments^16^.

**Figure 4.**
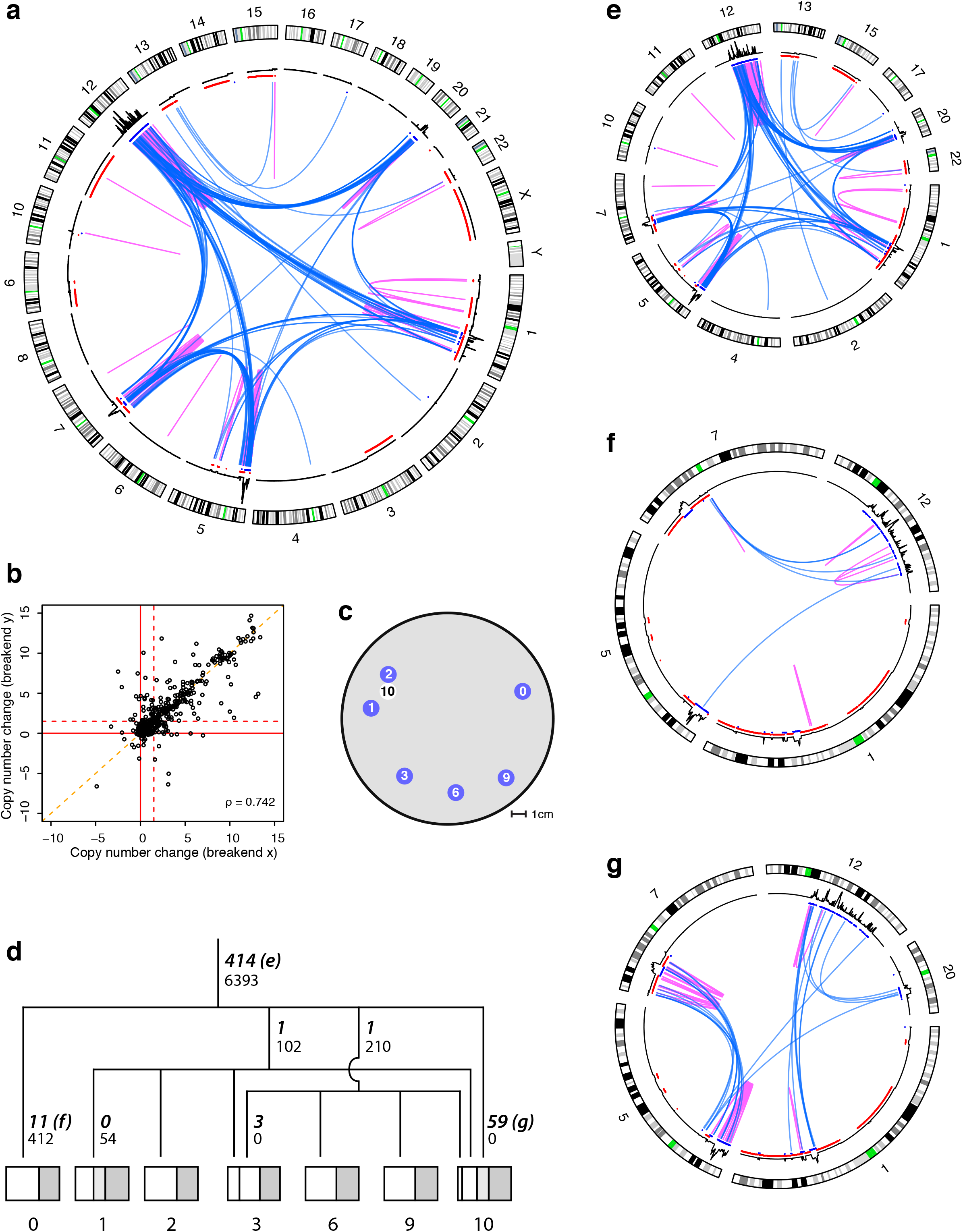
Somatic genome evolution of the sarcoma. (a), circos plot of all high-confidence breakpoint connections, indicated by arcs. Magenta, interchromosomal events; green, intrachromosomal. Otherwise, as in 3c. (b), scatterplot of copy number in the immediate vicinity of breakpoints. Each point is a breakpoint consisting of two breakends (X and Y, arbitrarily assigned) whose copy number estimates are plotted against each other. (c), location of the sampling sites from the sarcoma; 0 to 9 are from one cross-section, 10 is from a cross section parallel to it and separated by 3 cm. (d) Lineage tree of the samples reconstructed from high-confidence somatic SNVs. Number of SNVs supporting each branch are in small font, number of breakpoints are in bold italic with circos plot panel letters indicated for plots e-g. Samples are subdivided proportionally to somatic allele frequencies to indicate subclone size. Portion corresponding to normal contribution (e.g., infiltrating lymphocytes) is in dark grey. (e) Circos plots of the 414 ancestral (trunkal) events (f), events private to sample 0 (g), events private to sample 10.

One expectation regarding the detection of SVs using 10x data is that its high physical coverage improves the signal-to-noise ratio compared to standard short-read SV detection approaches. The number of SV-supporting 10x fragments correlated highly (rho=0.89; Supplementary Fig. 4) with the number of supporting mate-pairs, although the signal for mate-pair data was generally higher due to higher physical coverage. We also found a good correlation (rho=0.71; Supplementary Fig. 4b) between 10x and short-fragment support. Strikingly, there was a median 3.2 times as many 10x barcodes as short fragments supporting an event. The overall rate of validation of our breakpoints by mate-pairs was 94.6% (424/448), and this increased to 98.6% (351/356) when examining only those events that we were able to assemble. To compare the read cloud approach to previous methods, we applied commonly used tools to our standard Illumina libraries to identify large-scale SVs. We found that only 65.1% (375/576) of the short fragment-called somatic events were validated by mate-pair data (Supplementary Fig. 5).

We identified 239 somatic breakpoints in the sarcoma with at least 20 phased clouds supporting each side of the breakpoint. Of these events, the vast majority (229 or 96%) were supported by only a single haplotype combination, which is expected because the probability of the same exact SV occurring at the same position on both haplotypes is vanishingly small. In contrast, systematic errors resulting from, for example, genome repetitiveness, should affect all haplotypes equally. Therefore, the high percentage of events supported by only a single haplotype combination not only supports the validity of our phasing across breakpoints but also provides evidence that the breakpoint calls themselves do not result from substantial systematic biases.

### Genome evolution within the sarcoma

The 414 breakpoints present in all sarcoma samples but not in the control arose before the last common ancestor of the samples' cells. These shared, ancestral events include the chromothripsis on chromosome 12, with the vast majority of the other events involving chromosomes 1, 5, 7 and 20. In addition, we found an ancestral rearrangement followed by high-level amplification harboring the characteristic liposarcoma driver gene, *MDM2* (ref. 17).

We also identified 89 SVs that were present in certain subsets of the samples (but not in the control). The majority of these involved chromosomes 5, 7, and 12, and were private to one of the samples, marking subclone expansions that did not extend to the other samples: 59 in sample 10, 12 in sample 0 and 3 in sample 3. 3 had confounding copy number expansions in one or more samples, and 11 were clearly positive in sample 10 but exhibited very weak signal in a range of other samples, possibly also due to confounding copy number variation. Only 2 non-ancestral breakpoints were definitively shared by several samples and absent from the others.

The non-ancestral SVs and the inferred presence of subclones suggests that there was some evolutionary differentiation within the sarcoma that was captured by our sampling. We therefore set out to determine the evolutionary relationships amongst the samples and then analyze the dynamics of SV accumulation, based on the inferred phylogenetic tree. Because, as a class, single nucleotide variants (SNVs) are much more common than SVs, we turned to the short-fragment data to identify somatic SNVs and then build the samples' evolutionary tree based on the subset of phylogenetically informative SNVs^18^. In agreement with the SVs and copy number profiles, the majority (6393/7171) of high-confidence somatic SNVs were ancestral, originating before the last common ancestor of the samples’ cells. Four additional classes of SNVs were present in subsets of samples: one each that was private to samples 0 and 1, and two phylogenetically informative classes; these match the abovementioned sample combinations that harbored a single SV each.

The alternate allele frequencies of the SNVs of the two phylogenetically informative classes are highly consistent with the allele frequencies of the ancestral SNVs. The frequencies of SNVs present in the mixed lineage samples (3 and 10) are consistent with one another, with their sums matching the ancestral frequencies. The mutation spectrum of the somatic SNVs (data not shown) closely matches that of germline events, suggesting that they were caused by replication errors without special mutational mechanisms, and that they accumulated at a rate proportional to the number of cell divisions. Finally, as expected, the most phylogenetically similar samples were in close spatial proximity to one another within the tumor (Figure 4c). These lines of evidence support the idea that we were able to construct a robust evolutionary tree of our samples that could form the basis for interpreting the accumulation of SVs in this tumor (Figure 4d).

Analysis of the SVs on the basis of the tree suggests that SVs do not accumulate proportionally to the number of cell divisions and that they instead tend to occur in bursts, clustering in evolutionary time. Four branches in the tree are specifically informative in this regard: The two lineages that define the subclones of samples (1, 2, 3, 10) and (3, 6, 9) each only have one SV (2 out of 503) but a much larger proportion of SNVs (312 out of 7171). The private lineage of sample 10 has no SNVs (i.e., 0 out of 7171) but 59 breakpoints (out of 503). The private lineage of sample 0 has 412 out of 7171 SNVs, but only 11 out of 503 breakpoints; by contrast, the private lineage of sample 1 has 54 SNVs and 0 SVs. This utter lack of agreement between SNV and SV rates suggests that SV accumulation is episodic.

Further evidence for the episodic nature of SV accumulation is found in the differential localization of the breakpoints depending on exactly when they occurred during the evolution of the sarcoma. The 414 trunkal events are highly enriched for involvement of chromosome 12, mostly intrachromosomally, with some involvement of chromosomes 1, 5, 7 and 20 (Figure 4e). The private events in sample 1 mostly fell near regions of chromosomes 7 and 12 that harbor trunkal structural variation (Figure 4f). Strikingly, a large majority (43/59, 73%) of breakpoints present in the subclone private to sample 10 occurred within or between chromosomes 5 and 7 (Figure 4g). In contrast, only 30% of ancestral mutations occurred within or between those chromosomes. This enrichment was highly significant (p < 10^−9^, Fisher exact test), supporting the occurrence of a sudden series of events affecting a small portion of the tumor genome. These structural events thus likely occurred in a short enough time span that SNVs could not accumulate to substantial enough levels to directly observe the subclone.

## Discussion

One of the last frontiers in whole-genome sequencing, and in particular cancer genomics, is the accurate identification and reconstruction of complex structural variation^8^. Here we present a novel approach, GROC-SVs, that leverages read cloud information as implemented by 10x to discover structural variation. Applying it to multiple samples from a sarcoma and a breast cancer cell line we demonstrate the power of read clouds to not only dramatically reduce false discovery rates compared to standard short-fragment sequencing, but also enable the reconstruction of complex SVs.

To perform validation of the GROC-SVs breakpoint calls in the sarcoma, we used long-insert mate-pair libraries, the currently favored approach for detection of long distance genomic rearrangements and translocations^5,9,10^. 95% of somatic events were corroborated by the mate-pair data. In addition to enabling breakpoint detection, the read clouds allowed us to perform sequence assembly across breakpoints. Sequence-assembled SV calls had a 99% validation rate by the mate-pair data. In HCC1143, for which mate pair libraries were not available, we used sequence assembly of the linked reads involved in the breakpoint as a surrogate for validation of the breakpoint; 114/140 (81%) of somatic events could be successfully assembled.

Compared to read cloud data, standard short-fragment libraries provide lower physical coverage and lack long-distance information. In our data, SVs were typically supported by 3-fold more read clouds than fragments in the standard Illumina libraries. Thus, because of this lower breakpoint coverage, only 65% of somatic SVs identified from short-fragment libraries were validated by the mate pair data. Direct assembly of complex events was altogether impossible due to the lack of long-distance information.

To-date, genome-scale reconstruction of complex SVs has been limited to cases where the breakpoints are spaced no longer than the fragment insert size (typically ~500bp), or has involved indirect inference that events are related, based on their proximity and orientation in the reference genome^2,19^. Previous work has attempted to “walk” along chromothripsis events, finding pairs of breakpoints opposite one another and separated by a region of a single copy number^20,21^. One such analysis was conducted in a tour-de-force study of liposarcomas^16^, where neochromosomes were flow-sorted to improve sequencing coverage of these chromothripsis regions in standard Illumina libraries. In general, the accuracy of this painstaking walking process depends on the evenness of coverage in order to identify regions with similar copy number, and the sensitivity and accuracy of the method of detecting SVs.

Using the 10x data, we were able to directly reconstruct the order of large scale genomic rearrangements involving many breakpoints without the need for any assumptions about pairs of breakpoints. In the sarcoma genome, where chromothripsis produced dramatic genomic change, we found that 40% of our breakpoints fell within complex SVs, with adjacent breakpoints frequently separated by tens of kb. Notably, compared to prior approaches we accomplished this work without any difficult molecular biology in isolating material or building libraries, highlighting the practical potential of the 10x technology for characterizing large numbers of complex tumor genomes at a reasonable cost.

Most SVs in the sarcoma were shared across all 7 spatially distinct locations, and therefore must have occurred early in the evolution of the tumor. These ancestral events include the 174 chromosome-12 chromothripsis breakpoints and subsequent copy number amplifications as well an additional 240 breakpoints. In contrast, while 778 subclonal SNVs were detected, corresponding to 5 distinct subclone lineages, very few SVs other than the ancestral ones were shared across subclones. Thus, the sarcoma must have undergone an initial period of substantial structural instability, accumulating hundreds of rearrangements and copy number changes, before converging to a stable genomic configuration. This process appears to be similar to the liposarcoma chromothripsis followed by breakage-fusion-bridge cycles and subsequent chromosome linearization described by^16^.

Based on in vitro data, it has been hypothesized that copy number amplifications create a more permissive environment for chromothripsis since loss of the intervening segments would then result in the normal diploid, rather than haploid, copy number^22^. Consistent with a triploidy event preceding chromothripsis, but in contrast to most previous observations of chromothripsis, we found genomic segments falling between chromothriptic regions to be diploid, present at normal copy number, and to display both germline alleles of heterozygous single nucleotide polymorphisms.

In addition to the ancestral SVs, we found a small subclone private to sample 10 with 59 breakpoints that likely occurred in an additional, recent period of genome instability. These mutations largely occurred within and between chromosomes 5 and 7. We speculate that the ancestral SVs affecting chromosomes 5 and 7, including copy number amplifications, provided a more permissive environment for these private SVs to occur in, similar to the inferred ancestral triploidy event preceding chromothripsis of chromosome 12. Identification of this subclone despite a lack of detected SNVs within the clone highlights the substantial benefit of using SVs in addition to SNVs in understanding the evolutionary processes within a tumor.

In summary, using GROC-SVs, which we specifically developed for leveraging read cloud information, we show that 10x data allows for direct, data-driven reconstruction of complex structural variation. This is accomplished at high sensitivity and excellent specificity compared to short-fragment data, and at much lower laboratory effort and sample requirements than specialized libraries or mate pair approaches. Two distinct substrates, a chromothriptic sarcoma and a less highly rearranged breast cancer cell line, demonstrate wide applicability of the approach. Our evolutionary analysis of the sarcoma foreshadows substantial future advances in the related pursuits of reconstructing the full cancer genome and understanding each tumor’s structural evolution.

## Methods

### Sample Preparation and Library Construction

Sections (0.5cm thick, 14cm diameter) of a well-differentiated liposarcoma tumor, obtained under informed consent from the Stanford Tissue Bank, were cut into multiple pieces, snap frozen with liquid nitrogen, and stored at –80°C. Genomic DNA was extracted from 7 spatially distinct sites of this sarcoma as well as from matched control kidney tissue of the same patient. We extracted genomic DNA from about 20 mg tissue using Gentra Puregene Tissue Kit (Qiagen, Cat 158667). Tissue was ground in liquid nitrogen, lysed in Cell Lysis Solution and Proteinase K, and digested with RNase A. Protein was pelleted and removed by adding Protein Precipitation Solution followed by centrifugation. Genomic DNA was precipitated with isopropanol and resuspended in buffer EB). Purified genomic DNA was aliquoted and stored at –20°C.

Genomic DNA was separated by running about 1*µ*g DNA on a 1% low-melting-point agarose gel using Pulsed Field Gel Electrophoresis (PFGE). DNA of size 50–100 kb was then recovered by β-agarase I digestion and filter concentration (NEB, Cat M0392S). The size-selected DNA molecules of 1.2 ng were partitioned and barcoded using the 10x Genomics GemCode platform ^15^. Libraries were then sequenced with a HiSeq2500 to ~25-fold sequence coverage.

For short-fragment DNA libraries, 1 *µ*g of total genomic DNA was sheared to 350 bp. PCR-free libraries were then constructed using Illumina’s TruSeq DNA PCR-Free library preparation kit and sequenced with the Illumina HiSeqX system to ~35-fold sequence coverage.

For large-insert mate-pair libraries, 4 *µ*g of total genomic DNA was fragmented with Tagment Enzyme and gel size-selected to build 7kb-insert mate-pair libraries using Illumina’s Nextera Mate Pair Sample Preparation Kit (FC-132-1001) (Tagmentation, Strand Displacement, Gel Size Selection, Circularization, Linear DNA Digestion, Circulated DNA Shearing, Streptavidin Bead Binding, End Repairing, A-Tailing, Adaptor Ligation, and PCR Amplification). Libraries were sequenced with HiSeq2500 to ~20-fold sequence coverage.

### Breakpoint Detection

GROC-SVs is implemented as a multi-sample analysis pipeline, allowing the simultaneous analysis of multiple tumor and matched normal samples, or multiple related individuals.

GROC-SVs uses read alignments and (optionally) phasing information produced by the “Long Ranger” software from 10x Genomics. GROC-SVs begins by identifying all barcodes overlapping each 10 kb genomic window and then performing an all-by-all comparison. A pair of loci (x,y) is considered a structural variant candidate if the number of shared barcodes exceeds that expected based on the number of barcodes in each locus. For computational efficiency, this initial test is performed as a binomial test (a more rigorous test is applied later for each structural variant).

Next, candidate SV loci are clustered, and candidate breakpoints are extracted based on peaks in the distribution of read cloud ends. This takes advantage of the fact that read clouds are expected to end suddenly near each of the breakpoints; performing this operation only on those barcodes that are shared between the two loci dramatically improves both the signal and reduces the background. Candidate breakpoints are identified in each sample separately.

At this point, the breakpoints have been identified typically to within several kb of the correct location. The next step is to perform refinement on the breakpoint coordinates to obtain approximately nucleotide-level accuracy. This step takes all read clouds within 20 kb of the candidate site and selects only those clouds with barcodes shared on both sides of the breakpoint. Then, for each breakend (the two half-open intervals that make up each breakpoint) separately, the maximum point of read cloud density is found, and then walked toward the putative breakpoint location until the read cloud density drops off suddenly to background levels, indicating the presence of the breakpoint location. We found that this procedure typically identifies the correct breakpoint location to within several nucleotides if the breakpoint is uniquely mappable with short reads. In the case that the breakpoint region is not uniquely mappable, the inferred breakpoint location will be the last well-mappable (mapq ≥ 30) position before the breakpoint. Breakpoint refinement occurs across samples together so all fragments spanning a breakpoint are used for refinement, even if the event is only present in a small subclone within a sample.

### Sequence assembly of breakpoints

Next, a permissive clustering step groups breakpoints together if they share a substantial proportion of their barcodes. This is formulated as a simple threshold using the Jaccard Index, defined as the number of barcodes shared between the loci divided by the total number of barcodes. This Jaccard Index can be viewed as a sort of “allele frequency,” where the numerator counts the number of fragments supporting the event, and the denominator counts the number of fragments in the reference and alternate alleles. This is however an approximation because it is difficult to confidently assign any individual fragment to one allele since both reference- and alternate-allele-supporting fragments can end near either breakpoint location. Theoretically, another confounder is the non-zero rate of “barcode collisions”, where one fragment occurs near breakpoint x and an independent fragment occurs near breakpoint y, both in the same barcode. However, barcode collisions typically contribute a negligible amount to the numerator since the average number of barcode collisions is very small for most genomic regions (< 1 for GemCode and <<1 for Chromium in normal copy number regions, and only appreciably higher for extreme copy number outliers).

Within each cluster, the barcodes supporting each event are pooled together, and all reads originating from these supporting barcodes are collected. Sequence assembly is then performed on the collected reads using idba_ud^23^. As with breakpoint refinement, sequence assembly is performed multi-sample, so spanning fragments can be used for assembly even if they occur in samples with very low allele frequency. idba_ud was selected because its good performance across a wide range of sequence coverage, which is highest near the breakpoints and then low farther away. Contigs are then aligned against the reference genome and breakpoint locations are called where appropriate. Note that this assembly process may discover additional breakpoints that were not significant in the genome-wide breakpoint detection step for various reasons.

### Genome-wide reconstruction of complex events

Following sequence assembly, a more rigorous complex event reconstruction is performed. First, breakends sharing a substantial proportion of barcodes are again clustered together. The resulting clusters are represented as graphs with breakends represented as nodes, and connections between nearby (contiguous genomic segments) and distant (non-contiguous structural variants) breakends represented as edges. Because fragments may span many breakpoints at once, there may be barcode similarity between breakends that are separated by one or more breakpoints. Thus, for each breakend, we first select the assembly-supported breakpoint if one exists. The remaining breakpoints are selected based on the highest barcode support (nearby breakends should share more barcodes than distant ones). This process uses the high-quality information present in the sequence assemblies but can still perform complex event reconstruction even for breakpoints that cannot be sequence-assembled.

### Post-processing

During post-processing, a more rigorous p-value is assigned to each breakpoint. This p-value is calculated by randomly sampling the correct number of barcodes for each breakend from the background distribution of fragments per barcode, then calculating the number of shared barcodes. Resampling is performed 100 times, then the significance of the observed vs resampled number of shared barcodes is calculated using a ranksum test. This resampling procedure takes into account the effect of differences in genome coverage as well as the non-uniform partitioning of fragments across barcodes.

Additional filters are applied, primarily for use when analyzing germline events to identify candidate segmental duplications (segmental duplications should be present in both tumor and matched normal samples and are thus removed when analyzing somatic events). One filter of note compares the observed fragment lengths across breakpoints to those expected based on the background distribution. Structural variants should show long fragment support at 10s of kb away from each breakend. In contrast, segmental duplications and other repetitive genomic sequences often result in short supporting read clouds.

A final post-processing step assigns a present/absent call to each event for each sample. This genotype combines the resampling p-value calculated above as well as requiring a minimum allele frequency (again calculated using the Jaccard Index). Note that heterozygous and homozygous calls are not calculated because these are difficult to accurately define for the different types of structural variant and especially when copy numbers are variable.

### Validation

Mate-pair validation was performed by counting the number of mate-pairs in the expected orientation and distance relative to the two breakends. We used only reads with a very conservative mapping quality filter of mapq ≥ 55. The rationale for this high mapq filter was that true events should typically have mates mapping several kb away from the breakend, escaping any local repetitiveness around a breakend. We analyzed the background distribution of random genomic regions, and found that the vast majority of regions shared zero mate-pairs, and thus we used a conservative cutoff of 50 mate-pairs to consider an event to be validated. We also tried a more lenient cutoff of 10 mate-pairs with similar results.

### Evolutionary analysis

Evolutionary trees relating samples within the sarcoma were built as in refs 18 and 24.

Copy numbers were not used in the detection of SVs and were only calculated to gain a better understanding of the context for SVs. Because the coverage profiles for the 1^st^ generation 10x GemCode libraries showed substantial GC bias, we used standard PCR-free Illumina libraries to calculate copy number, normalized to the matched normal and normalized for DNA content within a sample. Coverage was typically higher for the tumor samples because of the many, large single-copy genomic regions.

## Acknowledgements

We thank Kristina Giorda, Sofia Kyriazopoulou-Panagiotopoulou, and Michael Schnall-Levin for their assistance in preparing and analyzing the 10x data, and Daniele Ramazzotti for analyzing mutation spectra. This work was supported by the Stanford Center for Computational, Evolutionary and Human Genomics (NS), R01CA183904 (NHI/NCI; RBW, SB, AS), and the BRCA Foundation (AS). Certain commercial equipment, instruments, or materials are identified in this document. Such identification does not imply recommendation or endorsement by the National Institute of Standards and Technology, nor does it imply that the products identified are necessarily the best available for the purpose. GROC-SVs is open source and available at https://github.com/grocsvs/grocsvs. Sequencing data are being submitted to the appropriate archives and will be made publicly available prior to publication of this work.

## Author Contributions

NS, ZW, AB, RBW, JMZ, MS, SB and AS designed the experiments and/or analyses. NS, ZW, JM and DC conducted the experiments. NS wrote analysis software. NS and AS analyzed the data. NS and AS wrote the manuscript with input from all authors.

